# Mechanisms of far-red light-mediated dampening of defense against *Botrytis cinerea* in tomato leaves

**DOI:** 10.1101/2021.01.21.427668

**Authors:** Sarah Courbier, Basten L. Snoek, Kaisa Kajala, Saskia C.M. Van Wees, Ronald Pierik

## Abstract

Plants detect neighboring competitors through a decrease in the ratio between red and far-red light (R:FR). This decreased R:FR is perceived by phytochrome photoreceptors and triggers shade avoidance responses such as shoot elongation and upward leaf movement (hyponasty). In addition to promoting elongation growth, low R:FR perception enhances plant susceptibility to pathogens: the growth-defense trade-off. Although increased susceptibility in low R:FR has been studied for over a decade, the associated timing of molecular events is still unknown. Here, we studied the chronology of FR-induced susceptibility events in tomato plants pre-exposed to either white light (WL) or WL supplemented with FR light (WL+FR) prior to inoculation with the necrotrophic fungus *Botrytis cinerea* (*B.c.*). We monitored the leaf transcriptional changes over a 30-hr time course upon infection and followed up with functional studies to identify mechanisms. We found that FR-induced susceptibility in tomato is linked to a general dampening of *B.c.-*responsive gene expression, and a delay in both pathogen recognition and jasmonic acid-mediated defense gene expression. In addition, we found that the supplemental FR-induced ethylene emissions affect plant immune responses under WL+FR conditions. This study increases our understanding of the growth-immunity trade-off, while simultaneously providing leads to improve tomato resistance against pathogens in dense cropping systems.

**One-sentence summary:** Low Red:Far-red ratio enhances tomato susceptibility towards the necrotrophic fungus *Botrytis cinerea* via delayed early pathogen detection and dampening of jasmonic acid-mediated defense activation.

## Introduction

Plants need sufficient light capture to sustain their photoautotrophic growth. At high planting densities, leaves grow closely and even overlap. This results in reduced light availability and changes in light quality due to preferential absorption of red (R, 600-700nm) and blue light (B, 400-500nm) and the reflection of far-red light (FR, ̴750nm) towards neighboring plants in the canopy. Changes in the red: far-red ratio (R:FR) are sensed by a specialized family of photo-convertible photoreceptors known as phytochromes where phytochrome B (phyB) plays the major role in *Arabidopsis thaliana* (Arabidopsis) (Franklin, 2008). A decrease of R:FR promotes the conversion of phyB from its active (Pfr) into its inactive (Pr) form. The photoinhibition of phyB by FR-enriched light leads to the release of PIF (Phytochrome-Interacting Factors) transcription factors, which subsequently bind G and E boxes in the promoters of target genes to initiate shade avoidance-associated gene expression (Franklin, 2008; Lorrain et al., 2008; Franklin and Quail, 2010; Casal, 2013). PIF target genes are associated with auxin homeostasis and cell wall remodeling to control growth (Hornitschek et al., 2012; Li et al., 2012; Pedmale et al., 2016). Also, auxin biosynthesis and transport allows for coordination of growth responses across an organ or even the entire organism in response to heterogeneous light conditions (Kohnen et al., 2016; Pantazopoulou et al., 2017; Küpers et al., 2018). Low R:FR also promotes gibberellin (GA) biosynthesis via the induction of the GA biosynthesis genes *GA20ox* and *GA3ox* (reviewed in Ballaré and Pierik, 2017). Upon GA binding to GIBBERELLIN INSENSITIVE DWARF1 (GID1) receptors, it initiates the ubiquitination and subsequent degradation of the negative growth regulators, DELLAs. Together, this results in PIF-mediated growth responses associated with rapid hypocotyl and petiole elongation as well as increased leaf angles (hyponasty) in Arabidopsis, characterized as the shade avoidance syndrome (SAS) (Franklin, 2008; Casal, 2012).

Plant responses to (a)biotic stresses are also affected by low R:FR (Ballaré and Pierik, 2017; Courbier and Pierik, 2019). In Arabidopsis, additional FR radiation has been shown to enhance plant susceptibility towards an array of pathogens showing a strong interplay between light and defense signaling pathways. Low R.FR conditions negatively affect both salicylic acid and jasmonic acid (JA)-responsive gene expression, which are crucial hormonal pathways that induce downstream defense responses towards plant resistance (De Wit et al., 2013). Interplay between JA and the growth promoting hormone GA plays a role in compromised plant resistance in low R:FR conditions and this occurs, at least partly, via interaction between growth-inhibiting DELLA proteins and defense-suppressing JASMONATE-ZIM DOMAIN (JAZ) proteins. The low R:FR-induced increase in bioactive GA leads to reduced DELLA levels, thus releasing JAZ proteins that can suppress defense (Hou et al., 2010; Cerrudo et al., 2012; Yang et al., 2012). Low R:FR-induced downregulation of JA-mediated defense is further associated with an increased stability of JAZ10 (Leone et al., 2014; Cerrudo et al., 2017), and a decrease in bioactive JA levels in Arabidopsis (Fernández-Milmanda et al., 2020). These observations that low R:FR perception lead to growth promotion at the expense of defense responses is known as the “growth-defense tradeoff”, which was originally thought to results from passive sink-source interactions and is now known to involve fine-tuned molecular control mechanisms in the plant (Campos et al., 2016; Ballaré and Austin, 2019; Major et al., 2020).

Light-dependent modulation of growth and susceptibility has mainly been studied in Arabidopsis, but has sporadically been studied in other species, including tomato. Low R:FR reduces tomato defenses against chewing and piercing insects (Izaguirre et al., 2006) and tomato *phyB1phyB2* double mutants exhibit increased leaf damage caused by *Mamestra brassicae* caterpillars compared to wild type plants (Cortés et al., 2016). Interestingly, also indirect defense are promoted by phyB inactivation in tomato: a changed JA-regulated pool of volatile organic compounds attracted more *Macrolophus pygmaeus* predatory insects (Cortés et al., 2016). We observed recently that supplemental FR does not only repress resistance against herbivorous insects, but also against the necrotrophic pathogen *Botrytis cinerea* (Ji et al., 2019; Courbier et al., 2020). In fruiting tomato plants, supplemental FR has been shown to enhance growth, fruit set and dry mass partitioning towards tomato fruits while it also promoted foliar *Botrytis cinerea* (*B. cinerea*) lesion development (Ji et al., 2019). We also recently demonstrated that supplemental FR lead to an increase in soluble sugars in tomato leaves responsible for increased lesion development induced by *B. cinerea* (Courbier et al., 2020b). Nevertheless, little is known still about the timing of events involved in the supplemental FR-pathogen interaction and the associated transcriptome reprogramming.

*B. cinerea* is one of the most destructive pathogens worldwide (Dean et al., 2012) and various transcriptome analyses have been performed on the Arabidopsis-*B. cinerea* pathosystem on both host and pathogen (Windram et al., 2012; Zhang et al., 2019; Soltis et al., 2020). Here, we aim to unravel the effect of supplemental FR on the timing of hormonal and metabolic pathway transcriptional induction during *B. cinerea* infection of susceptible tomato plants that are often cultivated in dense stands. We performed a transcriptome analysis of a 30 hr infection time course following a FR pretreatment which, to our knowledge, is the first time series experiment describing temporal changes of tomato in response to *B. cinerea* after a supplemental FR exposure. Our data show that supplemental FR pretreatment leads a delay in *B. cinerea*-induced defense activation, by altering JA and possibly ethylene-dependent pathways. The transcriptome analysis highlighted a set of six *PROTEINASE INHIBITOR* genes (*PI*) that are induced only in WL-treated samples. Using these likely defense-associated genes as markers, we further resolve the roles of JA and ethylene in FR-induced susceptibility in tomato.

## Results

### FR-enriched light promotes shoot elongation and susceptibility towards *B. cinerea*

To investigate the morphological changes induced by FR light enrichment (WL+FR), four-weeks-old tomato plants were exposed to either WL or WL+FR for five days and several growth parameters were recorded daily. Upon WL+FR exposure, plants exhibit a strong stem elongation compared to WL conditions (Fig. 1a and 1b). Although petiole length also increased upon WL+FR exposure (Fig. 1c), plants did not show a hyponastic (upward leaf movement) response, a typical shade avoidance response in Arabidopsis (Pantazopoulou et al., 2017), but remained constant while the WL-treated plants exhibited some epinastic (downward) leaf movement over time (Fig. 1d), probably because of leaf aging. In addition, tomato plants showed an increased lamina area and dry weight but no difference in specific leaf area (SLA) (Fig. S2). Even though the leaf area increases slightly, the leaf thickness remained unchanged for leaves already formed before the start of the WL+FR treatment (Fig. S2a and Fig. 1e). We also investigated the effect of FR supplementation applied before and/or during interaction with *B. cinerea* (Fig. 2). The lesion area was clearly larger on plants pretreated with WL+FR before inoculation compared to WL-pretreated plants irrespective of the light treatment applied after inoculation (Fig. 2). Although we used a standard bioassay on excised leaflets, we obtained similar findings in intact, whole plant systems (Fig. S3). Larger lesion size was associated with an increase in *B. cinerea* genomic DNA content in WL+FR-treated leaf tissue showing a positive effect of FR supplementation on *B. cinerea* growth *in planta* (Fig. S4a). We confirmed this *in vitro* by showing that *B. cinerea* mycelium biomass was increased by approximately 30% by WL+FR, even though the mycelium diameter on these plates remained unchanged (Fig. S4b and S4c). Altogether, our results show that WL+FR exposure of tomato plants elicits strong tomato phenotype changes and enhanced performance of *B. cinerea* on these plants. As the outcome of the infection was not influenced by the light quality applied after inoculation, we conclude that a WL+FR pretreatment alters subsequent plant responses to *B. cinerea* and promote its development in infected plant tissue.

**Figure 1:**
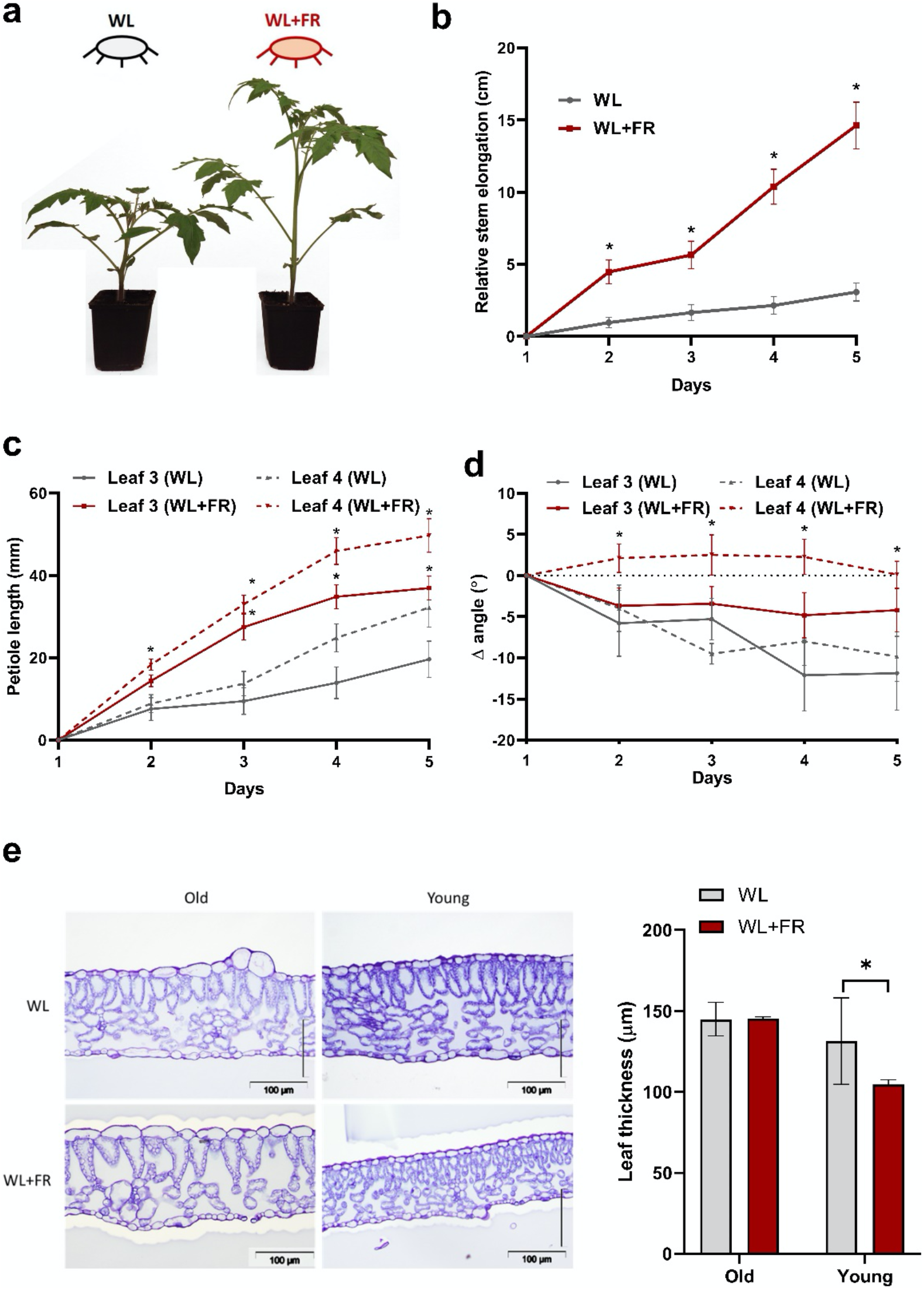
Supplemental FR light induces architectural changes in tomato plants. (**a**) Three-week-old tomato plants photographed after five days in WL or WL+FR conditions. (**b**) Stem elongation, (**c**) petiole length as well as (**d**) leaf angle of the third and fourth oldest leaf were recorded every day (ZT = 3). (**e**) Thickness measurements performed on tomato leaflets exposed to either WL or WL+FR for five days. Measurements were performed on leaflets that were either pre-existing (old) prior to the start of the light treatment or newly-formed (young) during the 5-day light treatment. Data show mean ± SEM and asterisks represent significant differences between WL and WL+FR-exposed plants (and for leaf 3 and leaf 4 independently (**c** and **d**)) according to Student’s t-test (p < 0.05), n = 4 – 10.

**Figure 2:**
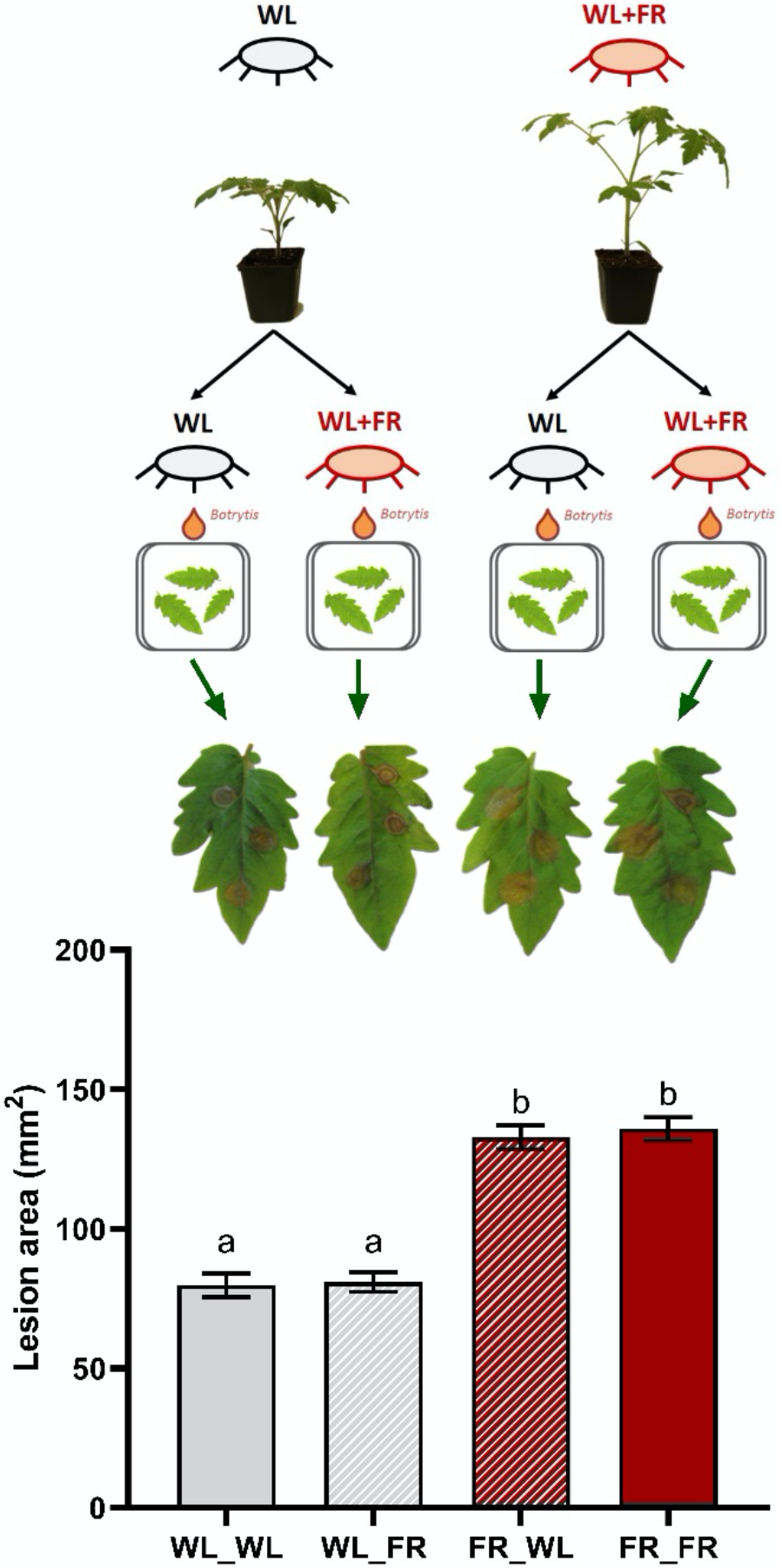
Supplemental FR light enhances tomato susceptibility towards *Botrytis cinerea*. Leaflets from the third oldest leaf of three-week-old tomato plants exposed to WL or WL+FR conditions for five days were detached, drop-inoculated with *B. cinerea* spores and incubated in either WL or WL+FR for 3 days. Plotted data represent the lesion area measured on the leaflets at 3 dpi (days post inoculation). Data show mean ± SEM and different letters represent significant differences between treatments according to ANOVA, Tukey’s post-hoc test (p < 0.05), n = 7 – 8 plants per treatment.

### Gene expression dynamics are affected by supplemental FR and *B. cinerea*

We aimed to understand the mechanisms underlying this WL+FR-induced susceptibility in tomato by investigating the associated transcriptome responses. Expression data were obtained from leaf tissue pre-exposed to WL or WL+FR prior to inoculation with *B. cinerea* spores or with a mock solution (Fig. 3a). During the infection, all detached leaflets were placed under WL conditions, excluding direct growth-promoting effects of FR on *B. cinerea* itself (Fig. S4b and S4c), and thus allowing us to study the plant responses specifically. The experiment consisted of 95 samples collected over two pretreatments (WL and WL+FR light), two treatments (mock versus infection with *B. cinerea* (*B.c.*)), six time points and three to four replicates per sample (Fig. 3a, Supplemental dataset S2). We first performed a principal coordinate analysis (PCoA) where the mapped reads segregated according to the light treatment at 6 and 12 hpi and the effect of the infection became visible at the later timepoint (24 hpi and 30 hpi) (Fig. 3b). These dynamics also reflected the number of differentially expressed genes (DEGs) where thousands of genes were modulated upon WL+FR pre-exposure at 6 and 12 hpi and upon infection at later timepoints (Fig. 3c-e). At 0 hpi, only few genes were modulated by WL+FR exposure (15 up- and 23 downregulated genes), implying that light quality does not extensively influence the tomato transcriptome after an extended period of time but rather upon reexposure to WL after inoculation (Fig. 3c). On the contrary, *B. cinerea*-responsive genes are modulated at later time points, likely resulting from a lag phase between the inoculation and the actual infection of leaf cells (Fig. 3d and 3e). Importantly, *B. cinerea*-responsive genes are detected from 12 hpi onwards in WL-pretreated samples while these are detected only from 24 hpi onwards in WL+FR-pretreated samples, suggesting a substantial delay in defense gene activation (Fig. 3d and 3e, respectively). In addition, many more genes were modulated by *B. cinerea* in WL-pretreated samples (6511 DEGs) compared to WL+FR-pretreated samples (2944 DEGs). Altogether, these observations indicate drastic FR-mediated dampening of *B. cinerea*-responsive transcriptome response.

**Figure 3:**
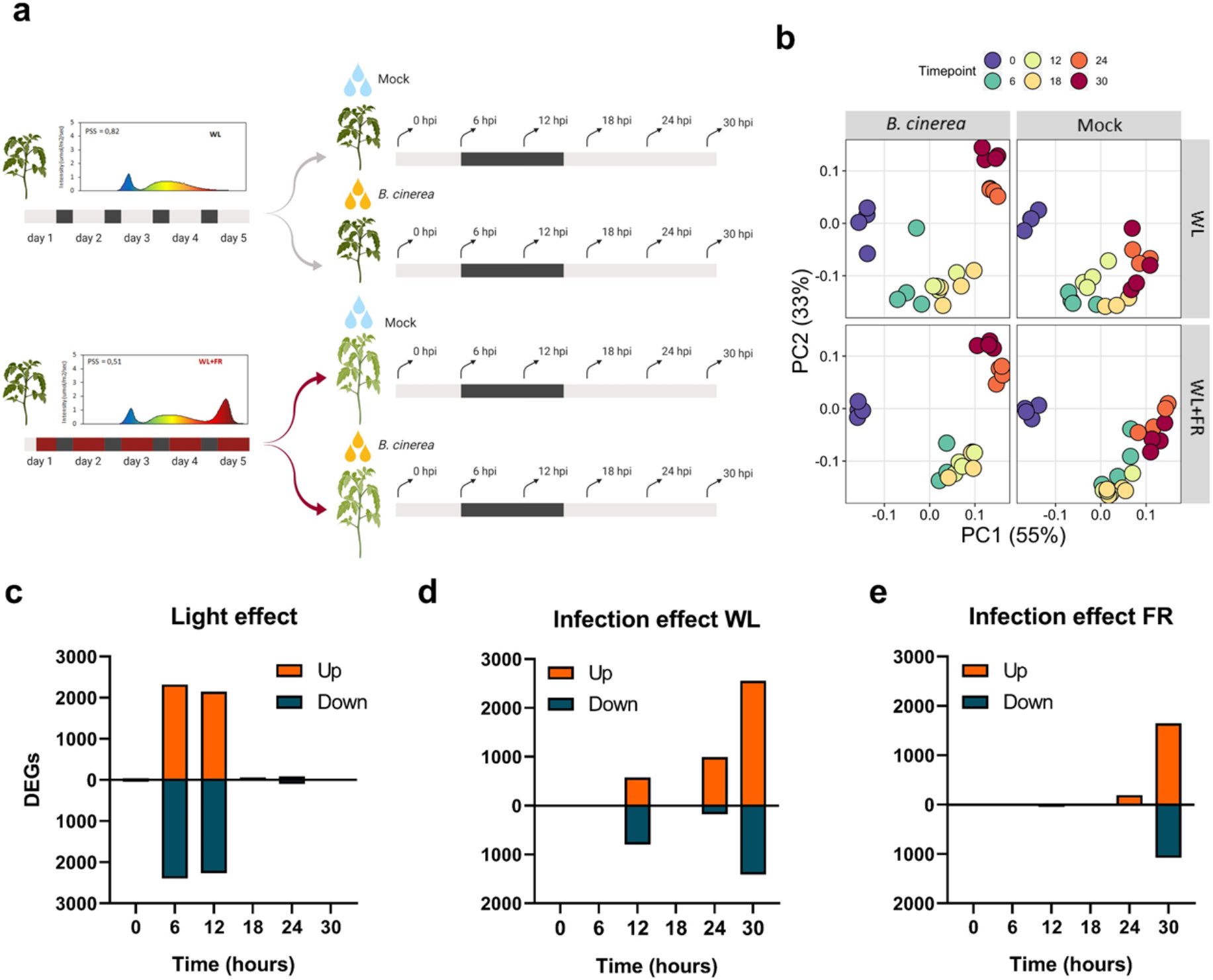
Responses to light and infection are not simultaneous and differ under WL+FR compared to WL conditions. **(a)** Experimental set up and harvest time points for the time series of *B. cinerea* infection on tomato leaflets of WL- or WL+FR-pretreated plants. Four weeks old tomato plants exposed for 5 days in WL (grey) or WL+FR (red) condition prior to drop-inoculation with *B. cinerea* or treatment with mock solution. Spectra represent the light background. The dark grey area represents an 8-h dark period. After 5 days of light pretreatment, leaflets were detached and drop-inoculated with *B. cinerea* spores or a mock solution as a control and harvested at 0, 6, 12, 18, 24 and 30 hpi (hours post inoculation) represented by the arrows. The infection was performed under WL conditions. (**b**) Visualization of the variation in gene expression by principal coordinate analysis (PCoA). The analysis was separated per treatment namely WL, WL+FR, *B. cinerea* and Mock conditions. Different colors represent the different timepoints and each dot corresponds to a replicate (0, 6, 12, 18, 24, 30 hpi). (**c-e**) Differentially expressed genes (DEGs) through time indicating upregulated (orange) and downregulated (dark blue) DEGs by comparing (**c**) mock-treated samples in WL+FR compared to WL, (**d**) *B. cinerea*-infected samples compared to mock samples after a WL or (**e**) WL+FR pretreatment based on BH method for multiple testing adjusted p value p <0.05.

### Light impacts gene expression at early timepoints

To investigate the direct effect of light quality on gene expression over time, we selected DEGs affected by WL+FR pre-exposure in the absence of the pathogen. Most of the supplemental FR-mediated gene expression changes were observed at 6 and 12 hr after leaf excission and return to WL (Fig. 3b and 3c). By performing a GO term enrichment on all WL+FR-responsive genes for each timepoint, we observed the upregulation of GO categories related with oxidoreduction processes, regulation of transcription, proteasome activity and glycolytic process, indicating an effect of WL+FR pre-exposure on gene expression and metabolism through time (Fig. 4a). At the same time we noticed a downregulation of for example glucan- and cell biogenesis-related processes upon WL+FR pretreatment at 12 hpi (Fig. 4b). The oxidoreduction GO term enrichment could hint at regulation of oxidative stress responses upon pathogen detection by supplemental FR and we, therefore, performed a luminol-based hydrogen peroxide (H2O2) quantification. Plants were elicited with the bacterial elicitor flagellin (flg22) to initiate a reactive oxygen species (ROS) burst. A strong and rapid increase in H2O2 production occurred upon elicitation with flg22 (Fig. 5a and 5b), but this burst was severely suppressed in WL+FR-pretreated leaf tissue (Fig. 5a). Also, WL+FR-pretreated plants displayed decreased basal H2O2 levels compared to WL (Fig. 5b) indicating that WL-pretreated plants might be better prepared to respond to a pathogen attack by producing ROS (Fig 5a and 5b). Based especially on the GO enrichments found for downregulated genes (Fig. 4b), we tested whether supplemental FR could affect cell wall and/or membrane integrity. By performing an electrolyte leakage assay, we observed an increase in the percentage of electrolyte loss in the WL+FR-pretreated plants compared to WL (Fig. 5c) which could in principle assist pathogen entry as WL+FR negatively affects membrane integrity.

**Figure 4:**
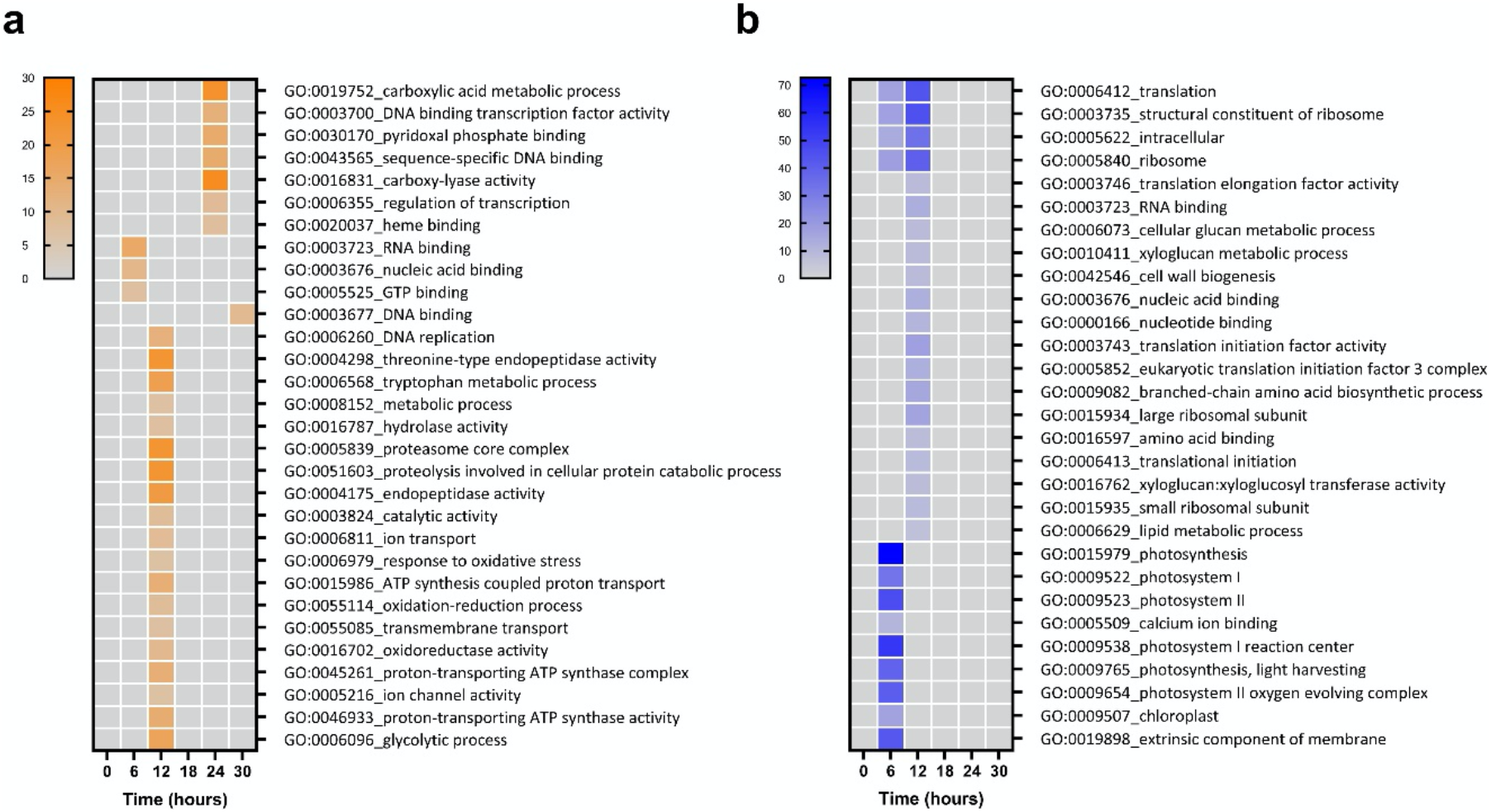
Gene ontology analysis of DEGs in WL+FR pretreatment compared to WL pretreatment under non-infected conditions. Heatmap of the gene ontology (GO) categories based on ‒log_10_ p-value where shades of orange (**a**) represent enrichment in upregulated DEGs and blue (**b**) in the downregulated DEGs.

**Figure 5:**
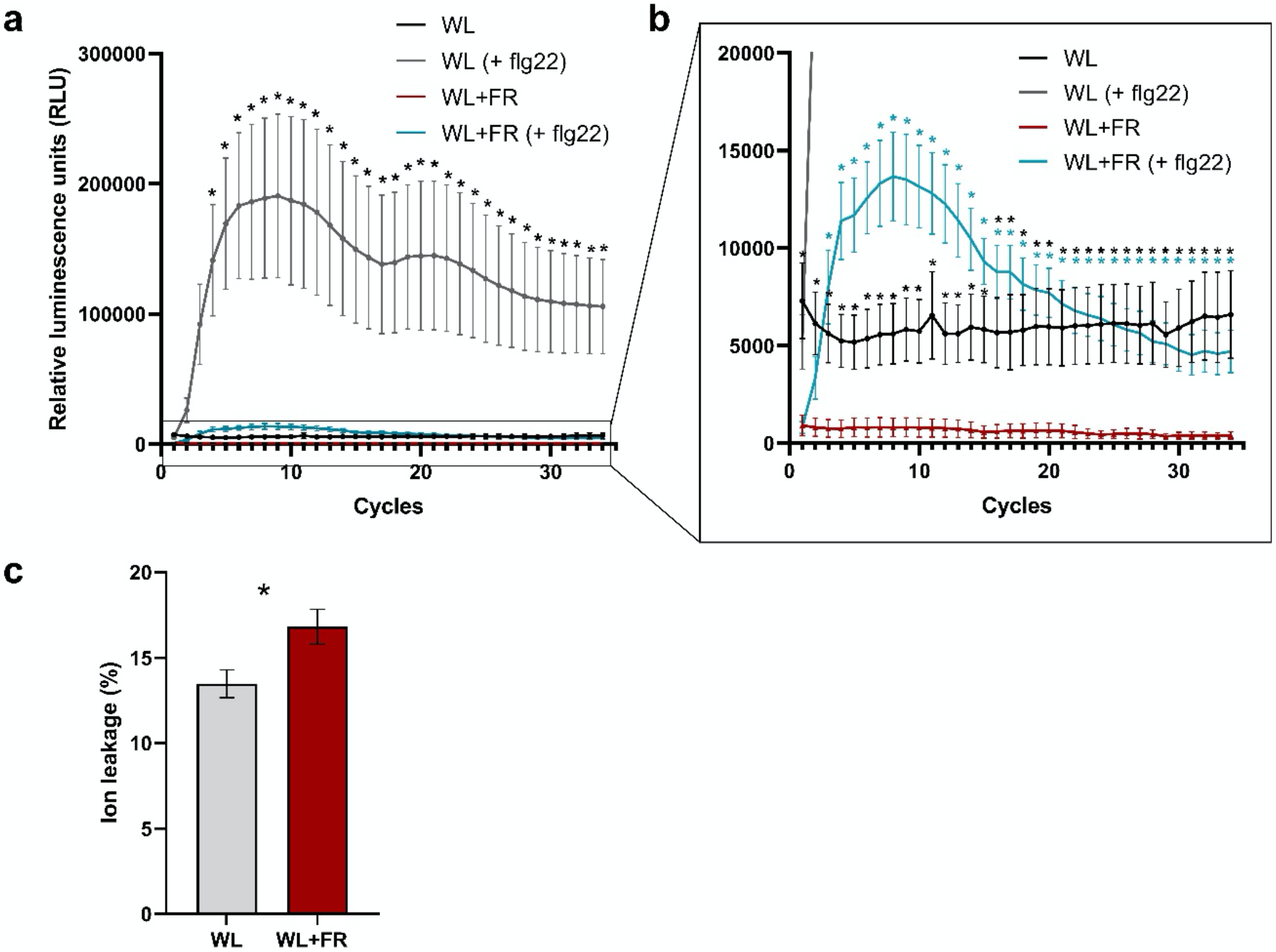
Supplemental FR pre-exposure alters ROS production and cell integrity. **(a and b)** ROS quantification by luminescence on tomato leaf discs originating from four-week old tomato plants treated for 5 days in WL or WL+FR light conditions. Leaf discs were exposed to either the bacterial elicitor flagellin (+ flg22) or were left unelicited and taken as controls. Data represent mean ± SEM and each measurement (cycle) lasted approximately 100 sec. n = 8. (**c**) Electrolyte leakage quantification on WL and WL+FR-treated leaf discs. Percentage of electrolytes loss from tomato leaf discs originating from four-week-old tomato plants exposed for five days to WL or WL+FR light conditions. Data represent mean ± SEM. Asterisk represents significant difference according to Student’s t-test (p < 0.05), n = 5.

### Supplemental FR light delays pathogen recognition and dampens defense activation

As *B. cinerea*-responsive gene modulation was delayed by WL+FR (Fig. 3d and 3e), we tested all *B. cinerea*-responsive genes for GO term enrichment (Fig. 6). Out of 76 enriched processes found upregulated upon *B. cinerea*-infection in WL-pretreated samples, only 34 were shared between WL and WL+FR-treated samples. From these 34 GO categories shared between both light pre-treatments, 14 categories were delayed by WL+FR light (Fig. 6a). At 12 hpi, we observed a striking inhibition of GO categories associated with protein catabolic processes (GO:0006511 and GO:0051603) via the proteasome (GO:0005839 and GO:0019773) or endopeptidase activity (GO:0004175, GO:0004190 and GO:0004298) by WL+FR (Fig. 6a). At later timepoints, we found enrichment of GO terms associated with chitin binding (GO:0008061) and chitin degradation processes (GO:0006032 and GO:0004568) in WL but not in WL+FR-pretreated samples. This could suggest that supplemental FR inhibits plant response to degrade the fungal cell wall. Consistently, genes associated with responses to biotic stimulus or defense response upregulated at 24 hpi in WL, are absent or delayed in WL+FR (Fig. 6a). These observations fit the notion that WL-pretreated plants recognize the pathogen and actively restrict its progression better than WL+FR-pretreated plants. At 24 and 30 hpi in WL+FR-pretreated samples, we also observed a delay in the upregulation of genes associated with protein phosphorylation (GO:0006468) and protein kinase activity (GO:0004674 and GO:0004672), possibly involved in downstream defense responses. DNA binding transcription factor activity (GO:0003700) was upregulated in WL-pretreated samples at 24 hpi but absent in WL+FR-pretreated samples. This category was composed of genes encoding WRKY transcription factors of which some might be involved in salicylic acid (SA) and jasmonic acid (JA) signaling (Pandey and Somssich, 2009) and Ethylene Responsive Factors (ERF) mainly regulated by ethylene (Müller and Munné-Bosch, 2015).

**Figure 6:**
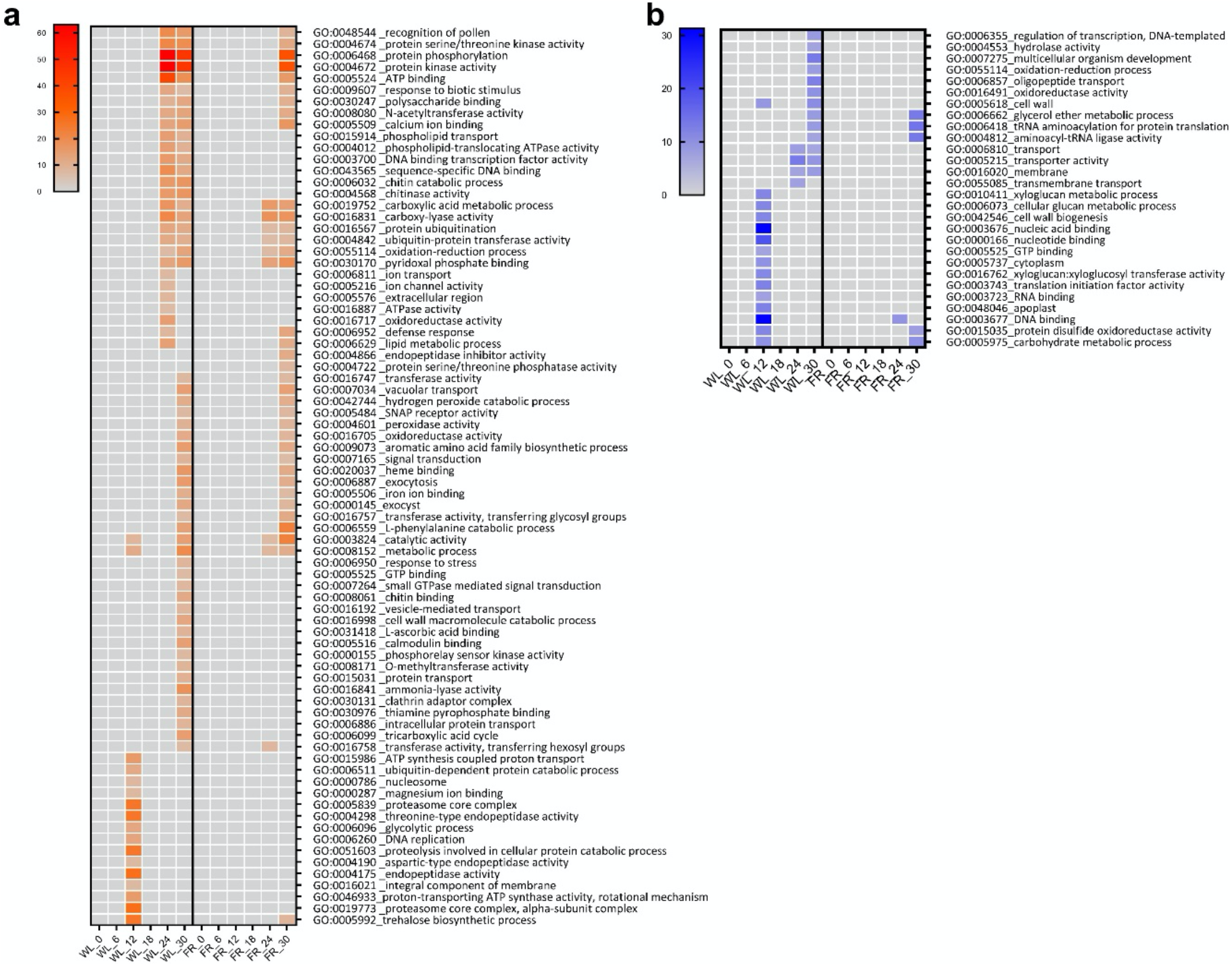
Gene ontology analysis of *B. cinerea*-responsive genes. Heatmap of the Gene ontology categories (GO) based on (–)log_10_(p-value) of *B. cinerea-*responsive DEGs in WL and WL+FR light exposed plants where orange (**a**) represent enrichment in upregulated DEGs and blue (**b**) in downregulated DEGs.

The involvement of ethylene was confirmed by an upregulation of ethylene biosynthesis genes *1-AMINOCYCLOPROPANE-1-CARBOXYLIC ACID* (ACC) *OXIDASE* and *SYNTHASE* (*ACO* and *ACS*) at 24 and 30 hpi in WL and only at 30 hpi in WL+FR-pretreated plants (Fig. 6a). Despite the delayed induction of ethylene biosynthesis genes upon infection with *B. cinerea* WL+FR-pretreated plants, ethylene emissions were elevated by WL+FR as compared to WL (Fig. 7); a classic effect of supplemental FR (Kegge and Pierik, 2010). Altogether, our data indicate that supplemental FR affects both ethylene and JA signaling in tomato defense against *B. cinerea*. Upon infection, only 6 out of 28 downregulated categories found in WL-pretreated samples were shared with WL+FR-pretreated samples (Fig. 6b). Interestingly, GO categories associated with xyloglucan and cellular glucan metabolism as well as xyloglucan:xyloglucosyl transferase activity and cell wall biogenesis (GO:0010411, GO:0006073, GO:0016762 and GO:0042546) were strongly downregulated at 12 hpi in WL conditions only (Fig. 6b). The downregulation of glucan-related structural processes could potentially reflect carbohydrates reallocation towards energy production (*e.g.* GO:0015986 and GO:0046933; Fig. 6a) to respond to the attacker. Interestingly, we did not observe such phenomenon in WL+FR-pretreated plants probably as the fungus was not yet detected at 12 hpi. Combined with previous research showing that WL+FR-pretreated plants display elevated soluble sugars levels in leaves promoting disease severity (Courbier et al., 2020), the delay in transcriptome changes in WL+FR-pretreated plants could be an additional explaination to the increased susceptibility of these plants. Taken together, our results indicate that *B. cinerea* infection triggers strong transcriptome reprogramming in WL which is significantly dampened, delayed, or non-existent in WL+FR-pretreated samples, and likely associated with increased susceptibility.

**Figure 7:**
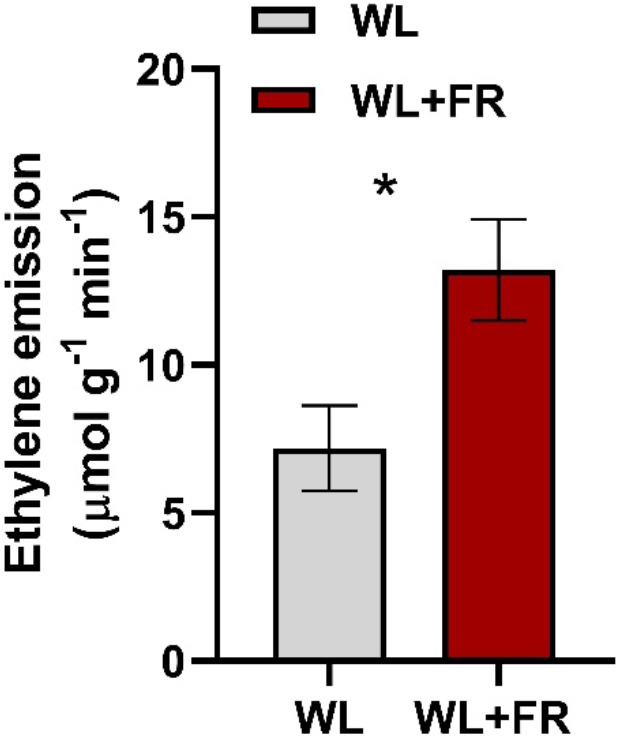
Leaf ethylene emissions are increase by supplemental FR. Ethylene emission quantified after five days of WL and WL+FR light treatments. n = 6-8. Data represent mean ± SEM. Asterisk represents significant difference according to Student’s t-test (p < 0.05).

### Supplemental FR-induced susceptibility coincides with downregulation of hormonal signaling

To define the core gene expression module associated with FR-induced susceptibility through time in tomato, we performed a 3-way ANOVA on light*infection*time to select genes interactively modulated by light, infection and time (Fig S5). We then ran a GO enrichment analysis on this group of genes and identified 11 significantly over-represented categories (Table 1, Supp. dataset S8). One of these categories was associated with response to wounding (GO:0009611) and was composed of six genes, all encoding PROTEINASE INHIBITORS (PI). These six genes were upregulated at 12 hpi upon *B. cinerea* infection in WL-pretreated samples only and not in the WL+FR-pretreated samples (Fig. 8a). *PI* genes are JA-responsive and highly induced upon *B. cinerea* infection in tomato (El Oirdi et al., 2011) indicating involvement of JA in the FR-induced susceptibility and a possible alteration of the JA-mediated *PI* gene induction by WL+FR (Fig. 8a). RPKM-normalized expression profiles for these six genes in all conditions tested in the RNA-seq data showed that all *PI* genes seem to be transiently upregulated in response to *B. cinerea* in WL conditions while this induction was strongly dampened in WL+FR conditions (Fig. 8b-g). Interestingly, we observed a peak of expression of all genes at 6 hpi in WL-treated samples in the absence of *B. cinerea* possibly reflecting the wounding responses triggered by detaching the leaflets from the plants prior to the bioassays. In WL+FR conditions, this putative wound-mediated *PI* peak was either delayed (Fig. 8b-d and 8g) or absent (Fig. 8e and 8f). Altogether, these results show that *PI* genes are responsive to *B. cinerea* and possibly wounding and their expression is dampened and/or delayed by WL+FR.

**Table 1:**
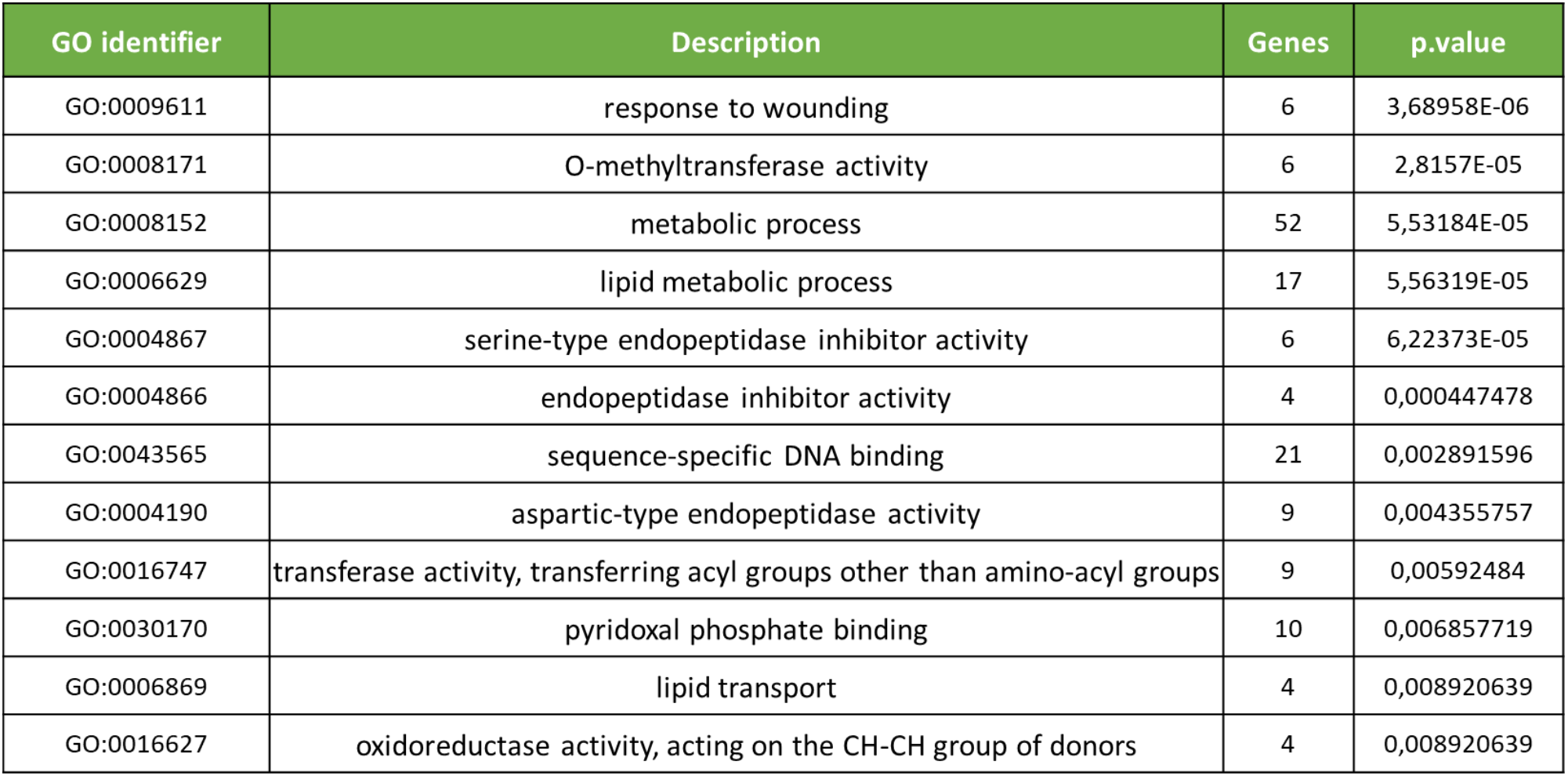
Gene ontology categories enriched in DEGs from the light*infection*time interaction.

| GO identifier | Description | Genes | p.value |
| --- | --- | --- | --- |
| GO:0009611 | response to wounding | 6 | 3,68958E-06 |
| GO:0008171 | O-methyltransferase activity | 6 | 2,8157E-05 |
| GO:0008152 | metabolic process | 52 | 5,53184E-05 |
| GO:0006629 | lipid metabolic process | 17 | 5,56319E-05 |
| GO:0004867 | serine-type endopeptidase inhibitor activity | 6 | 6,22373E-05 |
| GO:0004866 | endopeptidase inhibitor activity | 4 | 0,000447478 |
| GO:0043565 | sequence-specific DNA binding | 21 | 0,002891596 |
| GO:0004190 | aspartic-type endopeptidase activity | 9 | 0,004355757 |
| GO:0016747 | transferase activity, transferring acyl groups other than amino-acyl groups | 9 | 0,00592484 |
| GO:0030170 | pyridoxal phosphate binding | 10 | 0,006857719 |
| GO:0006869 | lipid transport | 4 | 0,008920639 |
| GO:0016627 | oxidoreductase activity, acting on the CH-CH group of donors | 4 | 0,008920639 |

**Figure 8:**
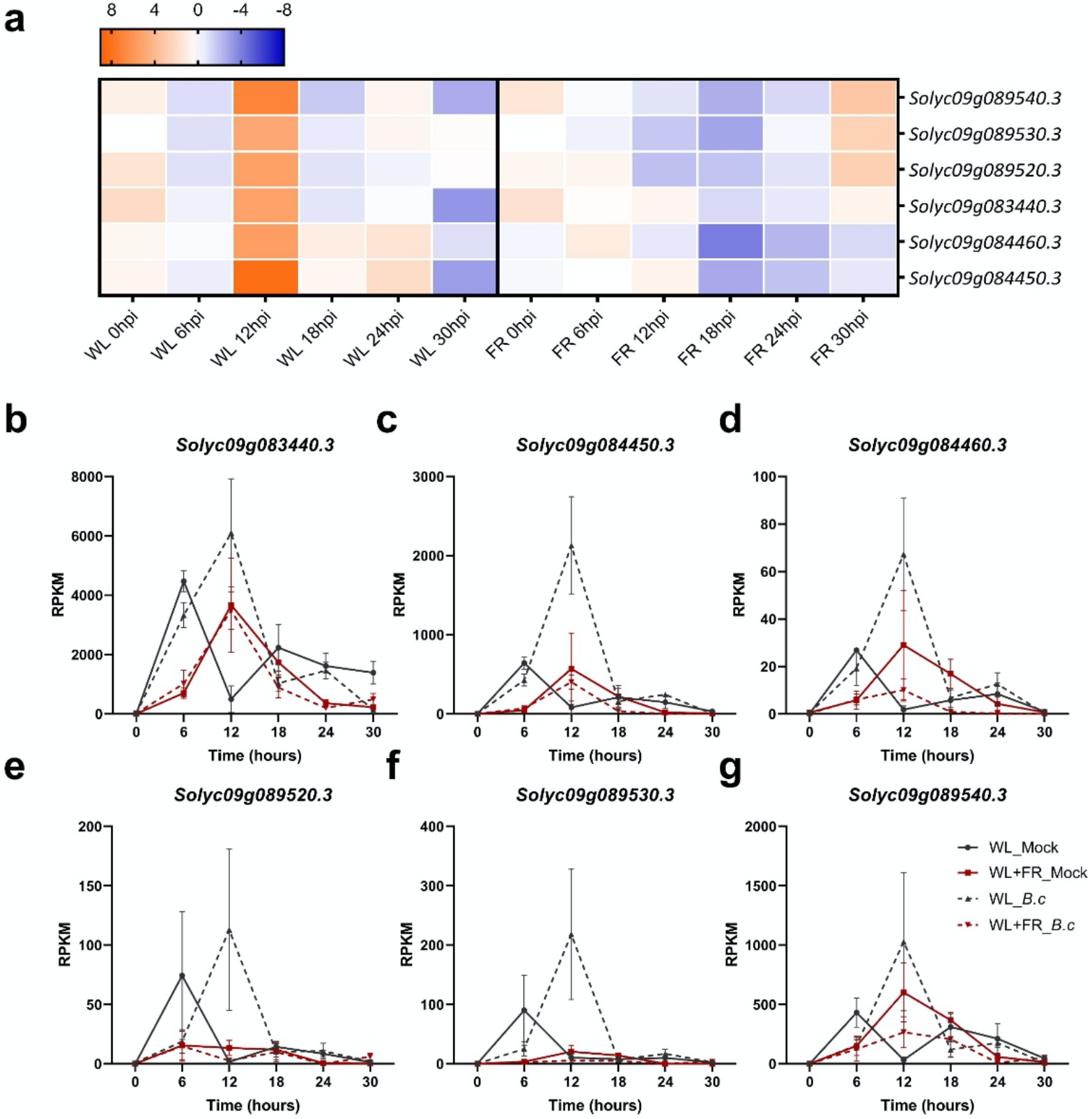
Dynamics of expression of *PROTEINASE INHIBITOR* (*PI*) genes significantly interacting in the 3-way light*infection*time ANOVA. **(a)** Heatmap corresponding to the log2FC of six *PI* genes in response to *B. cinerea* infection after a 5-day WL or WL+FR pretreatment at 0, 6, 12, 18, 24 and 30 hpi. Orange and blue colors represent up- and downregulation, respectively. (**b-g)** Expression patterns (RPKM) of the six *PI* genes. Each plot corresponds to the expression patterns of an individual gene through time in the four conditions tested, namely WL (grey) or WL+FR (red) pre-exposed plants inoculated with *B. cinerea* (dashed lines) or treated with a mock solution (plain lines). Plotted data represent the mean of replicates per timepoints ± SEM.

### Supplemental FR delays activation of jasmonic acid signaling

To confirm the trends of *PI* gene expression (Fig. 8b-g), and further study the involvement of JA in the regulation of these genes, we sprayed the 3^rd^ leaf of intact tomato plants with 100 μM MeJA or a mock solution and quantified mRNA abundance 15 min and 4 hr later (Fig 9). After 15 min, we observed a strong induction of *PI* genes under WL conditions, which was significantly reduced in WL+FR conditions for five out of the six *PI* genes (Fig 9a-d and 9f). After 4 hr, there was still a significant reduction due to WL+FR in MeJA-induced expression levels of two of the *PI* genes (Fig 9a and 9c) but for the other four *PI* genes the induction level by MeJA was comparable between WL- and WL+FR-pretreated samples (Fig 9b and 9d-f). This suggests a delay rather than an overall inhibition in the induction of *PI* genes by MeJA under WL+FR enrichment.

**Figure 9:**
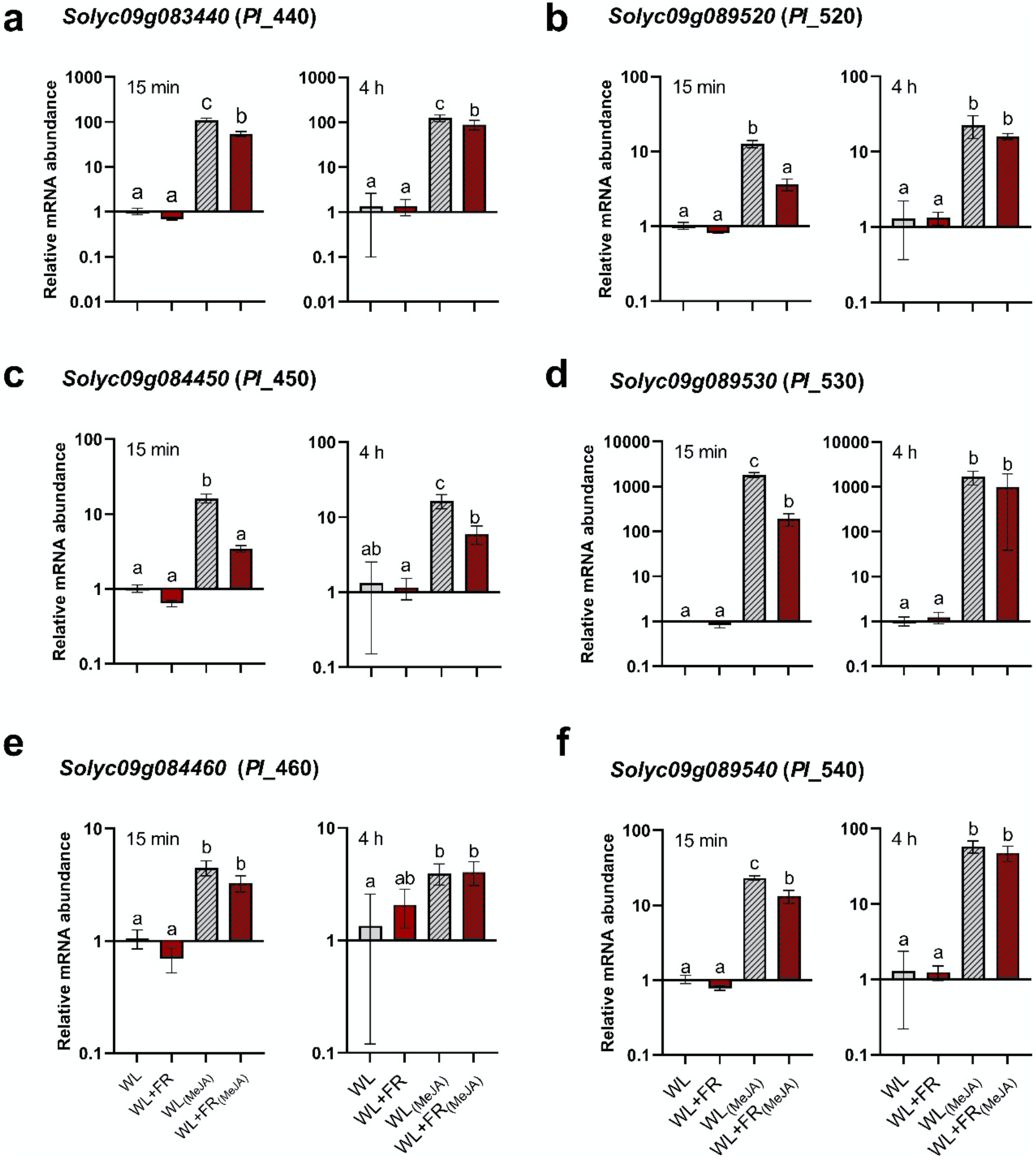
Supplemental FR causes a delay in JA-mediated response in tomato leaves. Transcript abundance analysis for six *PI* genes **(a-f)** after a WL (grey) or WL+FR (red) pretreatment followed by an exogenous MeJA treatment (100 μM; dashed bars) or a mock solution (plain bars) on intact plants. Leaf material was harvested at 15 min and 4 hours after the start of the MeJA treatment. Expression data are relative to WL conditions for each timepoint. Data represent mean ± SEM. Letters represent significant differences according to ANOVA, Tukey’s post-hoc test (p < 0.05).

### Supplemental FR interacts with JA and ethylene pathways to affect disease development

As defense responses against necrotrophic pathogens are mainly regulated by JA and ethylene (Penninckx et al., 1998), we assessed the involvement of these two hormones in tomato resistance towards *B. cinerea* and its modulation by supplemental FR. First, we tested the effect of exogenous MeJA (50 μM and 100 μM) and the JA biosynthesis inhibitor Jarin-1 (Fig. 10a and 10b). As expected, exogenous MeJA could enhance plant resistance in WL and this effect was stronger at increased MeJA concentrations confirming that JA is promoting defense (Fig. 10a). Although MeJA could also partly enhance the resistance in WL+FR-treated plants, the effect was only visible at the highest MeJA concentration applied, indicating that WL+FR-pretreated plants had reduced responsiveness to MeJA as compared to WL plants. The addition of Jarin-1 (50 μM) increased plant susceptibility to *B. cinerea* in WL-treated leaflets. However, it did not significantly affect the resistance of WL+FR-pretreated tissue as the lesion size upon Jarin-1 treatment was similar to the mock conditions (Fig. 10b). Altogether, these JA manipulations explain some of the WL+FR effects, but do not explain the full effect, implying other layers of resistance that can be altered by WL+FR. Since JA often acts together with ethylene, we also performed bioassays on detached leaflets treated with air (as a control), ACC (ethylene precursor), ethylene (C2H4), or 1-MCP (ethylene perception inhibitor) prior to inoculation with *B. cinerea* spores (Fig. 10c). Ethylene improved resistance in WL-treated plants, and ACC application improved resistance in both WL- and WL+FR-pretreated plant tissue. In addition, 1-MCP had minor effects in WL-pretreated leaflets while it further compromised tomato resistance in WL+FR-treated plants in line with the protective effect of ethylene in tomato resistance against *B. cinerea* (Díaz et al., 2002) (Fig. 10c). These data confirm that ethylene promotes resistance against *B. cinerea* in tomato, and indicate that the elevated ethylene emissions in WL+FR partly counterbalance the WL+FR-induced increased disease development upon *B. cinerea* infection.

**Figure 10:**
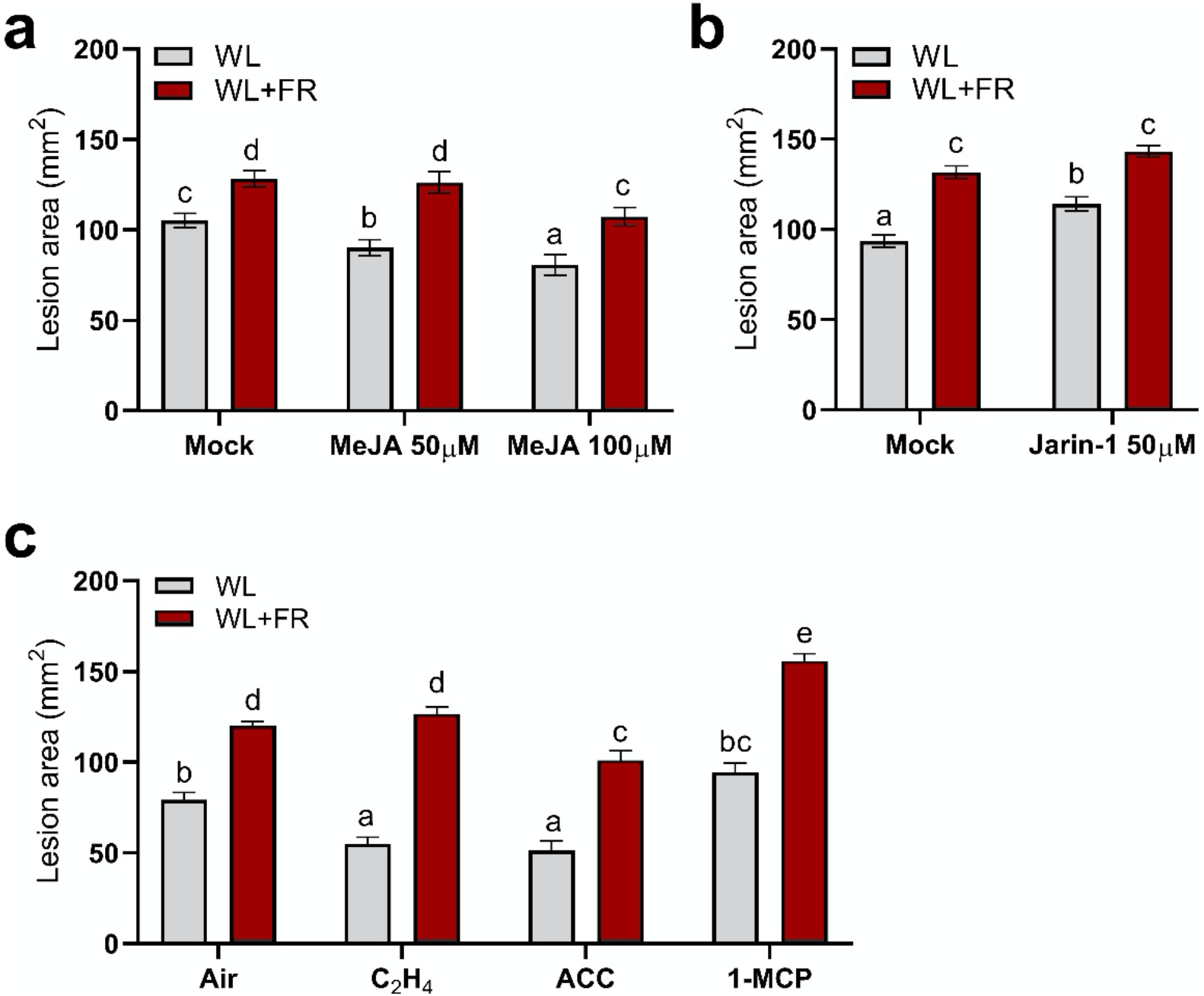
FR enrichment affects ethylene and JA-mediated defense in tomato towards *Botrytis cinerea.* Disease rating on tomato leaflets after 5 days of WL and WL+FR light treatments followed by **(a)** an exogenous MeJA (50 μM and 100 μM) or **(b)** Jarin-1 (JA biosynthesis inhibitor; 50 μM) or **(c)** exposure to either ethylene (C2H4), the ethylene precursor ACC or the ethylene receptor blocker 1-MCP treatment prior to inoculation in WL. n = 7-8 plants per treatment. Data represent mean ± SEM. Different letters represent significant differences according to ANOVA, Tukey’s post-hoc test (p < 0.05).

## Discussion

Here, we studied the chronology of tomato transcriptome events of WL and WL+FR-pretreated tomato plants upon infection with *B. cinerea* and unraveled multiple potential processes that could underlie the FR-induced susceptibility in tomato. Collectively, our findings suggest that WL+FR dampens metabolic, defense- and hormone-related processes involved in adequate plant defense against *B. cinerea* infection.

### Control of cell wall biogenesis and ROS-mediated plant defense by WL+FR

The cell wall is one of the first layers that pathogens encounter upon infection. The thickness and permeability of the cell wall and cell membrane might even be key to control pathogen penetration success (Underwood, 2012). In our conditions, we observed a downregulation of cell wall biogenesis upon FR exposure as well as upon *B. cinerea* infection and an increase in electrolyte leakage in WL+FR-pretreated plants compared to WL-pretreated plants (Fig. 4b, Fig. 5c and Fig. 6b) that could potentially facilitate pathogen penetration in plant tissue. Upon pathogen attack, the production of ROS by plants is essential for the establishment of resistance (Chinchilla et al., 2007) and we observed a clear upregulation of genes associated with oxidoreduction processes in WL+FR-pretreated samples compared to WL (Fig. 4a and Fig. 6a). Although genes associated with oxidoreduction processes are upregulated by supplemental FR, it is difficult to assess precisely whether WL+FR promotes oxidase or reductase activity. However, physiological data demonstrated that WL+FR-pretreated plants had a lower basal level of hydrogen peroxide (H_2_O_2_) and a much weaker H_2_O_2_ production burst upon elicitation with flagellin (Fig. 5a and 5b). It is possible that the strong upregulation of the genes associated with oxidoreduction processes could lead to a redox unbalance where plants cannot produce ROS as efficiently as in WL. However, we cannot exclude the possibility that WL+FR-pretreated plants over-scavenge ROS even in unstressed conditions resulting in reduced basal ROS production and weaker ROS-mediated defense responses.

### Supplemental FR delays pathogen detection and activation of defense signaling

Plants are able to recognize microbe-associated conserved motifs known as MAMPs at the plasma membrane as a very early signal preceding downstream defense gene activation (Jones and Dangl, 2006). Chitin is the main component of fungus cell wall and constitutes a very efficient MAMP that can trigger immediate defense gene activation upon a pathogen challenge (Zipfel, 2014). WL-pretreated plants showed an upregulation of chitin binding- and chitin degradation-associated processes upon *B. cinerea* infection (GO:0006032 and GO:0004568), whereas WL+FR-pretreated plants did not (Fig. 6a). Also, GO categories associated with “protein serine/threonine kinase activity”, “protein phosphorylation” and “protein kinase activity” that are enriched in upregulated DEGs at 24 hpi and 30 hpi in *B. cinerea*-challenged WL-pretreated plants are present only at 30 hpi in WL+FR-pretreated plants (Fig. 6a). Most of the DEGs associated with these three categories encode receptor like kinases (RLKs), leucine-rich repeats RLK (LRR-RLKs) and mitogen-activated protein kinases (MAPK). These are gene families of which members are often associated with pathogen detection at the plasma membrane and the subsequent defense gene activation (Zipfel, 2014). Interestingly, we also found *WALL-ASSOCATED_KINASE (WAK) -LIKE KINASE* (*Solyc02g090970*) upregulated in WL-treated plants only. As WAK1 has been identified as a galacturonides receptor and promoting *B. cinerea* resistance in Arabidopsis (Brutus et al., 2010), the lack of *Solyc02g090970* induction in supplemental FR would be consistent with the increased disease susceptibility and could indicate a putative delay in pathogen detection at the cell surface. This could in turn delay the induction of chitinases (Fig. 6a) that degrade the fungal cell wall, which would indirectly promote pathogen colonization in supplemental FR-pretreated leaf tissue compared to WL. In addition, a delayed pathogen detection at the cell surface correlates with a reduction of subsequent defense signaling, which is supported by a delayed representation of the GO categories associated to “defense response” and “response to biotic stimulus” in the supplemental FR-pretreated plants challenged with *B. cinerea* (Fig. 6a). Altogether, these observations indicate a clear effect of supplemental FR on response to infection, possibly due to a delay in pathogen recognition at the membrane that could result in a delay in chitin degradation and defense gene activation through the MAMP-mediated defense signalling cascade.

### Several JA-induced PI genes are suppressed by WL+FR

The 3-way ANOVA light*infection*time analysis revealed the core gene expression patterns associated with FR-induced susceptibility. Out of the 131 genes significant for the interaction, we found six *PROTEINASE INHIBITOR* (*PI*) genes highly induced upon infection by *B. cinerea* at 12 hpi in WL and not in WL+FR-pretreated samples (Fig. 8a and Fig. S4). *PI* genes have previously been reported to be strongly induced upon *B. cinerea* infection and promote plant resistance in a JA-dependent manner (El Oirdi et al., 2011). In our data, we observed that exogenous MeJA treatment induced *PI* genes within 15 min and that the level of expression was reduced in WL+FR-treated plants compared to WL. However, *PI* expression reached similar levels between WL and WL+FR-exposed plants after 4 hr of MeJA treatment showing either a reduced JA sensitivity or a delay in gene induction by supplemental FR-pretreatment (Fig. 9c-h). In addition, tomato resistance against *B. cinerea* was promoted by exogenous MeJA treatment while it was compromised by Jarin-1 in WL plants (Fig. 9a and 9b). However, we could only rescue resistance in WL+FR samples at the highest applied concentration of 100 μM MeJA while 50 μM already promoted resistance in WL-treated plants confirming the hypothesis of a lower JA sensitivity in WL+FR (Moreno et al., 2009; Cerrudo et al., 2012; Izaguirre et al., 2013; De Wit et al., 2013). As supplemental FR appears to reduce JA responsiveness (Fig. 9), it is likely that plants do not respond to JA-inducing *B. cinerea* infection as effectively as WL-exposed plants do, therefore allowing the pathogen to develop faster on leaves treated with supplemental FR (Fig. S3a). In addition, we showed that the susceptibility of WL+FR-pretreated leaflets was always higher than WL-treated leaflets, even upon high concentration of exogenous MeJA, pointing towards other layers of direct or indirect immunity. We showed previously that WL+FR-treated tomato leaves contain elevated soluble sugar levels, which promoted proliferation of *B. cinerea* (Courbier et al., 2020). Although this increased sugar accumulation was partially JA-dependent, it could not be fully explained by JA variations, and thus constitutes a partially independent pathway of supplemental FR-induced susceptibility (Courbier et al., 2020).

### Ethylene signaling is affected by supplemental FR

Among the GO processes and genes modulated upon infection, genes encoding ERF transcription factors were only upregulated in WL-pretreated samples and genes encoding ethylene biosynthesis enzymes were induced earlier in WL-pretreated plants compared to WL+FR upon infection with *B. cinerea* (Fig. 6a). We also observed that exogenous ACC or ethylene treatment promote plant resistance to the fungus (Díaz et al., 2002; Fig. 7). The promoting effect of ethylene alone on plant resistance was weak in WL+FR-pretreated plants, as compared to ACC. We expect that ACC continues to be converted into ethylene by the leaflets and therefore accumulates in the sealed petri dishes giving a long-lasting ethylene treatment compared to ethylene treatment alone, which is dissipated within minutes after opening the desiccators. Since ethylene is beneficial for plant defense, it is paradoxical that supplemental FR light would simultaneously promote ethylene emission, thus promoting resistance, while at the same time enhancing susceptibility through dampening of the JA pathway. Apparently, under low R:FR conditions, the elevated ethylene emissions exert some control over the enhanced susceptibility to pathogens. Indeed, when inhibiting the plant’s ethylene receptors with 1-MCP, disease development in the low R:FR-exposed plants became even more severe. We, therefore, conclude that under low R:FR conditions indicating proximate neighbors, plants downregulate the partially JA-dependent defense responses against *B. cinerea*, whilst upregulation ethylene emissions, which would tentatively help prevent an excessive proliferation of this pathogenic fungus.

## Conclusions

We demonstrated that plants experiencing WL+FR light display shoot and petiole elongation, typical shade avoidance traits accompanied by increased foliar susceptibility to *B. cinerea*. By performing a transcriptome analysis over a 30 h infection period, we demonstrate that supplemental FR strongly dampens and delays the expression of *B. cinerea*-responsive genes in tomato. We found that supplemental FR dampens the overall gene modulation in response to *B. cinerea* and especially JA-mediated responses in turn enhancing susceptibility in tomato. This phenomenon occurs via a reduced JA responsiveness postponing *PI* gene induction and the associated JA-mediated plant immune responses. In addtion, supplemental FR suppresses the classic ROS burst upon elicitor treatment and increases ion leakage. The elevated ethylene emissions in WL+FR-pretreated plants reduce pathogen development and may prevent excessive pathogen proliferation in low R:FR light conditions. The data collectively provide a broad overview and understanding of supplemental far-red-induced disease susceptibility. We anticipate this can aid further improvement of tomato genotypes for growth at high density.

## Materials and methods

### Plants growth conditions and light treatments

Tomato (*Solanum lycopersicum*) cv. Moneymaker (LA2706) seeds were obtained from the C. M. Rick Tomato Genetics Resource Center and propagated by the Horticulture and Product Physiology group at Wageningen University and Research as part of the LED it Be 50% consortium. Seeds were sown in wet vermiculite. After 10 days, seedlings were transferred into 9 x 9 cm pots with regular potting soil (Primasta® soil, the Netherlands). Tomato plants were grown for four weeks after sowing in climate chambers (MD1400; Snijders, The Netherlands) in long day photoperiod (8 hr dark / 16 hr light) at 150 μmol m^−2^ s^−1^ photosynthetically active radiation (PAR) using Philips GreenPower LED research modules (Signify B.V) white (WL, R:FR = 5.5) reaching a Phytochrome Stationary State (PSS) value of 0.8 (Sager et al., 1988). For supplemental FR treatments, three-week old plants were exposed for five days (starting at ZT = 3 on the first day) under WL supplemented with Philips GreenPower LED research modules far-red (FR) (WL+FR; R:FR = 0.14, PSS value = 0.5, PAR = 150 μmol m^−2^ s^−1^). The top two lateral leaflets of the 3rd leaf were used for all experiments. Light spectra used in this study were measured with a JAZ spectrophotometer (Ocean Optics Inc., UK) and are displayed in Fig. S1. Stem length were measured with a digital caliper. Petiole angles were measured from pictures with the ImageJ software. All measurements were calculated as day 5 – day 1.

### Leaf thickness measurements

Fresh leaf pieces were cut and immediately incubated in 1-2 ml of Karnovski’s fixative solution (2.1% formaldehyde, 2.5% glutaraldehyde and 0.1 M phosphate buffer (0.2 M NaH_2_PO_4_; 0.2 M Na_2_HPO_4_) at pH = 7). The samples were vacuum infiltrated for 10-15 min and slowly shaken at room temperature for 1 hr before three washes with MilliQ water. Samples were dehydrated by adding 1-2 ml of a graded series of ethanol (30, 50, 70, 90, 96%) replacing the ethanol every 30-60 min. Dehydrated leaf samples were embedded following the Technovit® 7100 plastic embedding system and shaped into a trapezoid with until the leaf tissue was visible in the middle. Sections of 10 μm were made using a Leica Om-U3 microtome and mounted on microscope slides supplemented with a drop of MilliQ water and the slides were dried on a heating plate (80 °C). The sections were stained with 0.5 % toluidine blue for 30-60 sec and washed thoroughly with MilliQ water. Images were taken and leaf thickness was measured using an Olympus fluorescent microscope BX50-WI.

### Fungal growth conditions and detached leaflets bioassays

*Botrytis cinerea* B05.10 was maintained on half strength potato dextrose agar medium (PDA ½, BD Difco™) and cultivated for two weeks under natural daylight conditions at room temperature. The spore suspension was prepared according to Van Wees et al. (2013) and diluted to a final concentration of 1.5 x 10^5^ spores ml^−1^ in half strength potato dextrose broth (PDB ½, BD Difco™) prior to the inoculation. Bioassays were performed on detached tomato leaflets previously treated in WL or WL+FR for five days. The leaflets were placed in square Petri dishes onto Whatman® filter soaked with 6 ml of tap water to avoid dehydration. The bioassays took place at 3 p.m. The adaxial side of the leaflets was drop-inoculated 3-6 times with 5 μl of spore suspension. Plates were sealed with PARAFILM® M and incubated for three days in their respective light treatment conditions (WL or WL+FR). Pictures were taken three days post inoculation (dpi) and lesion areas were measured using the imageJ software with the image processing package Fiji (Schindelin et al., 2012).

### ROS measurements

Tomato plants (four-week-old) were exposed to either WL or WL+FR for 4 days. On day 5, leaf discs (Ø 0.4 cm) originating from the 3^rd^ leaf of each plants were floated on deionized water for approximately 8 hr in either WL or WL+FR corresponding to the treatment light conditions at room temperature without shaking. Each leaf disc was placed in each well of a white flat-bottomed 96-well plate (Greiner LUMITRAC™ 200) onto 180 μl of deionized water mixed with 20 μl of 10X reaction mix (200 μM Luminol L-012 and 10 μg ml^−1^ horseradish peroxidase) with a final concentration of 20 μM Luminol and 1 μg ml^−1^ of peroxidase. The luminescence was quantified by using a GloMax luminometer (Promega). The background noise was measured for 15 min prior to adding 1 μM flg22 (final concentration) on the leaf discs. The luminescence was measured for approximately 1 hr by recording 34 cycles of 100 sec (Albert and Fürst, 2017).

### Ion leakage

Three leaf discs (Ø 0.6 cm) originating from the 3^rd^ oldest leaf of each plants were rinsed in deionized water to remove attached electrolytes leftover from the cutting. The conductivity was measured with an electrical conductivity meter after 3 hr of incubation in 10 ml of 400 mM Mannitol and again after 20 min at 95 °C. The electrolyte loss was calculated as a percentage of the total amount of electrolytes in leaf tissue indicated by the value measured after boiling.

### Chemical treatments

Five days after the start of the WL or WL+FR pretreatment, tomato leaflets from the 3^rd^ leaf were detached and immediately dipped for 10 sec in a 50 μM or 100 μM methyl-jasmonate (MeJA; Sigma-Aldrich) or mock solution (0.1 % EtOH) supplemented with 0.1 % Tween 20 then placed separate sealed Petri dishes. When performed on whole plants, the 3^rd^ leaf was sprayed with 100 μM MeJA or a mock solution (0.1 % Tween 20). Hormone treatments started at 10 a.m. in either WL or WL+FR. For gene expression experiments, leaf material was sampled for every condition at 0, 15 min and 4 hr after the start of the MeJA treatment. For bioassays, detached MeJA-treated tomato leaflets were inoculated in WL conditions with *B. cinerea* spores 4 hr after MeJA application. The same procedures and experiments were performed by using the JA biosynthesis inhibitor Jarin-1 (50 μM) (Meesters et al., 2014).

For ethylene-related experiments, tomato leaflets from the 3^rd^ leaf were detached and placed in square Petri dishes containing two discs of Whatman® filter paper soaked with tap water. Petri dishes were placed in separate transparent glass desiccators. Control plates were placed in a desiccator of which the lid was slightly opened to allow for air circulation. Ethylene treatments were performed by either replacing tap water by 6 ml of a 10 μM ACC (ethylene precursor) solution or by injecting gaseous ethylene (10 ppm) in the desiccator. Ethylene inhibitor treatments were performed by injecting 1-MCP (10 ppm) in the desiccator. All treatments lasted for 1 hr prior to inoculation with *B. cinerea* spores as previously described.

### Ethylene quantification by gas chromatography

Tomato leaflets were detached from WL- or WL+FR-pretreated plants, weighted, carefully rolled and inserted in a 1-ml syringe. After 30 min, the gas contained in the syringe was injected into a gas chromatography device (Syntech Spectras GC955) to determine ethylene levels.

### RNA isolation and gene expression analysis

Leaf discs treated with either JA or mock solution were sampled for RNA isolation (three or four biological replicate per treatment) and ground with silica beads for 1 min in a tissue lyser and supplemented with 300 μl of cell lysis buffer (2% SDS, 68 mM sodium citrate, 132 mM citric acid and 1 mM EDTA) and incubated for 5 min at room temperature prior to adding 100 ul of protein/DNA precipitation buffer (4 M NaCl, 16 mM sodium citrate and 32 mM citric acid) followed by 10 min of incubation on ice. Samples were centrifuged for 15 min at 13000 rpm, the supernatant was transferred in a new tube (300 μl) and supplemented with an equal volume of ice cold isopropanol. All tubes were centrifuged for 5 min at 13000 rpm, pellet was washed with 300 μl of 70% ethanol. RNA pellets were air dried for 5 min before elution in 30 μl RNAse-free water. cDNA synthesis was carried out using RevertAid H minus Reverse transcriptase (Thermo scientific). Gene expression analysis were performed by quantitative RT-qPCR in a Viia7PCR device with 5 μl reaction mix containing SyberGreen Supermix (Bio-Rad). Tomato actin was used as a reference gene and fold change in expression were calculated based on 2^^(-ddCt)^ (Livak and Schmittgen, 2001). Primer sequences used are displayed in Supp. data S1.

## Author contributions

R.P., S.W. and S.C. designed the research; S.C. carried out the biological experiments; B.L.S. performed the transcriptome analysis with input from K.K.; S.C., R.P. and K.K. discussed the results; S.C. and B.L.S. designed and made the figures; S.C wrote the manuscript with contributions of R.P., S.W., K.K. and B.L.S.

## Supplemental data

The following supplemental materials are available.

**Supplemental methods.** Methods used for supplemental data as well as for the RNA-sequencing time series bioassay, library preparation and sequencing data analysis.

**Supplemental figure S1.** LED light spectra used in this study.

**Supplemental figure S2.** Effect of supplemental FR on lamina area, lamina dry weight and specific leaf area.

**Supplemental figure S3.** Bioassay on whole plants exposed to WL or WL+FR.

**Supplemental figure S4.** qPCR on *B. cinerea* genomic DNA in infected leaf tissue and mycelium growth assay *in vitro*.

**Supplemental figure S5.** Heatmap corresponding to the log_2_FC for the 131 genes significant for the 3-way interaction (light*infection*time) in response to *B. cinerea* infection in either WL or WL+FR-pretreated plants.

**Supplemental dataset S1.** Primers used for conventional qPCR and qPCR on genomic DNA.

**Supplemental dataset S2.** RNA-sequencing sample description.

**Supplemental dataset S3.** RNA-sequencing raw read counts

**Supplemental dataset S4.** RNA-sequencing normalized read counts

**Supplemental dataset S5.** Correlation value between all samples

**Supplemental dataset S6.** Log_2_ fold change, p.value and adjusted p.value per gene, per comparison and per timepoints.

**Supplemental dataset S7.** Gene ontology (GO) enrichment per timepoint and per comparison.

**Supplemental dataset S8.** GO enrichment based on ANOVA (one, two or three factors).

The RNA sequencing materials are available on GEO public repository (accession number GSE157831, https://www.ncbi.nlm.nih.gov/geo/query/acc.cgi?acc=GSE157831).

## Acknowledgements

This work was funded by the Dutch Research Council (NWO) TTW Perspectief grant nr 14125 (LED it Be 50%) with contributions from Signify, LTO Glaskracht and WUR Greenhouse Horticulture. We thank the Utrecht Sequencing Facility for providing sequencing service and data. Utrecht Sequencing Facility is subsidized by the University Medical Center Utrecht, Hubrecht Institute, Utrecht University and The Netherlands X-omics Initiative (NWO project 184.034.019). We also thank UMC Utrecht Bioinformatics Expertise Core for data analysis and data handling. The UMC Utrecht Bioinformatics Expertise Core is subsidized by the University Medical Center Utrecht, Center for Molecular Medicine. We thank Tom Raaymakers for help with the ROS quantifications, Scott Hayes for help with the electrolyte leakage assays, Sjon Hartman for help with the ethylene experiments, Jesse Küpers and the LED it Be 50% consortium for helpful discussions.

## References

Albert M, Fürst U (2017) Quantitative detection of oxidative burst upon activation of plant receptor kinases. Methods Mol. Biol. Humana Press Inc. 69–76

Ballaré CL, Austin AT (2019) Recalculating growth and defense strategies under competition: key roles of photoreceptors and jasmonates. J Exp Bot 70: 3425–3434

Ballaré CL, Pierik R (2017) The shade-avoidance syndrome: Multiple signals and ecological consequences. Plant Cell Environ 40: 2530–2543

Brutus A, Sicilia F, Macone A, Cervone F, De Lorenzo G (2010) A domain swap approach reveals a role of the plant wall-associated kinase 1 (WAK1) as a receptor of oligogalacturonides. Proc Natl Acad Sci U S A 107: 9452–9457

Campos ML, Yoshida Y, Major IT, De Oliveira Ferreira D, Weraduwage SM, Froehlich JE, Johnson BF, Kramer DM, Jander G, Sharkey TD, et al (2016) Rewiring of jasmonate and phytochrome B signalling uncouples plant growth-defense tradeoffs. Nat Commun 7: 12570

Casal JJ (2013) Photoreceptor Signaling Networks in Plant Responses to Shade. Annu Rev Plant Biol 64: 403–427

Casal JJ (2012) Shade avoidance. Arabidopsis Book 10: e0157

Cerrudo I, Caliri-Ortiz ME, Keller MM, Degano ME, Demkura P V., Ballaré CL (2017) Exploring growth-defence trade-offs in Arabidopsis: phytochrome B inactivation requires JAZ10 to suppress plant immunity but not to trigger shade-avoidance responses. Plant Cell Environ 40: 635–644

Cerrudo I, Keller MM, Cargnel MD, Demkura P V, de Wit M, Patitucci MS, Pierik R, Pieterse CMJ, Ballaré CL (2012) Low red/far-red ratios reduce Arabidopsis resistance to Botrytis cinerea and jasmonate responses via a COI1-JAZ10-dependent, salicylic acid-independent mechanism. Plant Physiol 158: 2042–52

Chinchilla D, Zipfel C, Robatzek S, Kemmerling B, Nürnberger T, Jones JDG, Felix G, Boller T (2007) A flagellin-induced complex of the receptor FLS2 and BAK1 initiates plant defence. Nature 448: 497–500

Cortés LE, Weldegergis BT, Boccalandro HE, Dicke M, Ballaré CL (2016) Trading direct for indirect defense? Phytochrome B inactivation in tomato attenuates direct anti-herbivore defenses whilst enhancing volatile-mediated attraction of predators. New Phytol 212: 1057–1071

Courbier S, Grevink S, Sluijs E, Bonhomme PO, Kajala K, Van Wees SCM, Pierik R (2020) Far-red light promotes Botrytis cinerea disease development in tomato leaves via jasmonate-dependent modulation of soluble sugars. Plant Cell Environ 43:2769–2781

Courbier S, Pierik R (2019) Canopy light quality modulates stress responses in plants. iScience 22: 441–452

Dean R, Van Kan JAL, Pretorius ZA, Hammond-Kosack KE, Di Pietro A, Spanu PD, Rudd JJ, Dickman M, Kahmann R, Ellis J, et al (2012) The Top 10 fungal pathogens in molecular plant pathology. Mol Plant Pathol 13: 414–430

Díaz J, Ten Have A, Van Kan JAL (2002) The Role of Ethylene and Wound Signaling in Resistance of Tomato to Botrytis cinerea. Plant Physiol 129: 1341–1351

Fernández-Milmanda GL, Crocco CD, Reichelt M, Mazza CA, Köllner TG, Zhang T, Cargnel MD, Lichy MZ, Fiorucci AS, Fankhauser C, et al (2020) A light-dependent molecular link between competition cues and defence responses in plants. Nat Plants 6: 223–230

Franklin KA (2008) Shade avoidance. New Phytol 179: 930–944

Franklin KA, Quail PH (2010) Phytochrome functions in Arabidopsis development. J Exp Bot 61: 11–24

Hornitschek P, Kohnen M V., Lorrain S, Rougemont J, Ljung K, López-Vidriero I, Franco-Zorrilla JM, Solano R, Trevisan M, Pradervand S, et al (2012) Phytochrome interacting factors 4 and 5 control seedling growth in changing light conditions by directly controlling auxin signaling. Plant J 71: 699–711

Hou X, Lee LYC, Xia K, Yan Y, Yu H (2010) DELLAs Modulate Jasmonate Signaling via Competitive Binding to JAZs. Dev Cell 19: 884–894

Izaguirre MM, Mazza CA, Astigueta MS, Ciarla AM, Ballaré CL (2013) No time for candy: Passionfruit (Passiflora edulis) plants down-regulate damage-induced extra floral nectar production in response to light signals of competition. Oecologia 173: 213–221

Izaguirre MM, Mazza CA, Biondini M, Baldwin IT, Ballaré CL (2006) Remote sensing of future competitors: Impacts on plants defenses. Proc Natl Acad Sci U S A 103: 7170–7174

Ji Y, Ouzounis T, Courbier S, Kaiser E, Nguyen PT, Schouten HJ, Visser RGF, Pierik R, Marcelis LFM, Heuvelink E (2019) Far-red radiation increases dry mass partitioning to fruits but reduces Botrytis cinerea resistance in tomato. Environ Exp Bot. 168: 103889

Jones JDG, Dangl JL (2006) The plant immune system. Nature 444: 323–329

Kegge W, Pierik R (2010) Biogenic volatile organic compounds and plant competition. Trends Plant Sci 15: 126–132

Kohnen M V, Schmid-Siegert E, Trevisan M, Petrolati LA, Sénéchal F, Müller-Moulé P, Maloof J, Xenarios I, Fankhauser C (2016) Neighbor Detection Induces Organ-Specific Transcriptomes, Revealing Patterns Underlying Hypocotyl-Specific Growth. Plant Cell 28: 2889–2904

Küpers JJ, van Gelderen K, Pierik R (2018) Location Matters: Canopy Light Responses over Spatial Scales. Trends Plant Sci 23: 865–873

Leone M, Keller MM, Cerrudo I, Ballaré CL (2014) To grow or defend? Low red: Far-red ratios reduce jasmonate sensitivity in Arabidopsis seedlings by promoting DELLA degradation and increasing JAZ10 stability. New Phytol 204: 355–367

Li L, Ljung K, Breton G, Schmitz RJ, Pruneda-Paz J, Cowing-Zitron C, Cole BJ, Ivans LJ, Pedmale U V., Jung HS, et al (2012) Linking photoreceptor excitation to changes in plant architecture. Genes Dev 26: 785–790

Livak KJ, Schmittgen TD (2001) Analysis of relative gene expression data using real-time quantitative PCR and the 2-∆∆CT method. Methods 25: 402–408

Lorrain S, Allen T, Duek PD, Whitelam GC, Fankhauser C (2008) Phytochrome-mediated inhibition of shade avoidance involves degradation of growth-promoting bHLH transcription factors. Plant J 53: 312–323

Major IT, Guo Q, Zhai J, Kapali G, Kramer DM, Howea GA (2020) A phytochrome b-independent pathway restricts growth at high levels of jasmonate defense. Plant Physiol 183: 733–749

Meesters C, Mönig T, Oeljeklaus J, Krahn D, Westfall CS, Hause B, Jez JM, Kaiser M, Kombrink E (2014) A chemical inhibitor of jasmonate signaling targets JAR1 in Arabidopsis thaliana. Nat Chem Biol 10: 830–836

Moreno JE, Tao Y, Chory J, Ballare CL (2009) Ecological modulation of plant defense via phytochrome control of jasmonate sensitivity. Proc Natl Acad Sci 106: 4935–4940

Müller M, Munné-Bosch S (2015) Ethylene response factors: A key regulatory hub in hormone and stress signaling. Plant Physiol 169: 32–41

El Oirdi M, El Rahman TA, Rigano L, El Hadrami A, Rodriguez MC, Daayf F, Vojnov A, Bouarab K (2011) Botrytis cinerea manipulates the antagonistic effects between immune pathways to promote disease development in tomato. Plant Cell 23: 2405–2421

Pandey SP, Somssich IE (2009) The role of WRKY transcription factors in plant immunity. Plant Physiol 150: 1648–1655

Pantazopoulou CK, Bongers FJ, Küpers JJ, Reinen E, Das D, Evers JB, Anten NPR, Pierik R (2017) Neighbor detection at the leaf tip adaptively regulates upward leaf movement through spatial auxin dynamics. Proc Natl Acad Sci 114: 7450–7455

Pedmale U V, Huang SSC, Zander M, Cole BJ, Hetzel J, Ljung K, Reis PAB, Sridevi P, Nito K, Nery JR, et al (2016) Cryptochromes Interact Directly with PIFs to Control Plant Growth in Limiting Blue Light. Cell 164: 233–245

Penninckx IAMA, Thomma BPHJ, Buchala A, Métraux JP, Broekaert WF (1998) Concomitant activation of jasmonate and ethylene response pathways is required for induction of a plant defensin gene in Arabidopsis. Plant Cell 10: 2103–2113

Schindelin J, Arganda-Carreras I, Frise E, Kaynig V, Longair M, Pietzsch T, Preibisch S, Rueden C, Saalfeld S, Schmid B, et al (2012) Fiji: An open-source platform for biological-image analysis. Nat Methods 9: 676–682

Soltis NE, Caseys C, Zhang W, Corwin JA, Atwell S, Kliebenstein DJ (2020) Pathogen genetic control of transcriptome variation in the arabidopsis thaliana - Botrytis cinerea pathosystem. Genetics 215: 253–266

Underwood W (2012) The plant cell wall: A dynamic barrier against pathogen invasion. Front Plant Sci 3: 85

Van Wees SCM, Van Pelt JA, Bakker PAHM, Pieterse CMJ (2013) Bioassays for assessing jasmonate-dependent defenses triggered by pathogens, herbivorous insects, or beneficial rhizobacteria. Methods Mol Biol 1011: 35–49

Windram O, Madhou P, McHattie S, Hill C, Hickman R, Cooke E, Jenkins DJ, Penfold CA, Baxter L, Breeze E, et al (2012) Arabidopsis defense against Botrytis cinerea: chronology and regulation deciphered by high-resolution temporal transcriptomic analysis. Plant Cell 24: 3530–57

De Wit M, Spoel SH, Sanchez-Perez GF, Gommers CMM, Pieterse CMJ, Voesenek LACJ, Pierik R (2013) Perception of low red: Far-red ratio compromises both salicylic acid- and jasmonic acid-dependent pathogen defences in Arabidopsis. Plant J 75: 90–103

Yang DL, Yao J, Mei CS, Tong XH, Zeng LJ, Li Q, Xiao LT, Sun TP, Li J, Deng XW, et al (2012) Plant hormone jasmonate prioritizes defense over growth by interfering with gibberellin signaling cascade. Proc Natl Acad Sci U S A 109: E1192–E1200

Zhang W, Corwin JA, Copeland DH, Feusier J, Eshbaugh R, Cook DE, Atwell S, Kliebenstein DJ (2019) Plant–necrotroph co-transcriptome networks illuminate a metabolic battlefield. Elife. 8: e44279

Zipfel C (2014) Plant pattern-recognition receptors. Trends Immunol 35: 345–351

